# Inhibition of the neurodevelopmental disorder-associated 16p11.2 gene *QPRT* leads to altered cell type distribution in human stem cell-derived cerebral organoids

**DOI:** 10.1101/2025.09.08.673916

**Authors:** Julia Schwarzpaul, Clara M. Droell, Afsheen Kumar, Harishny Sarma, Madeleine Gruenauer, Selen Z. Ucar, Julio C. Hechavarría, Andreas G. Chiocchetti, Denise Haslinger

**Affiliations:** Goethe University Frankfurt, Department of Child and Adolescent Psychiatry, Psychosomatics and Psychotherapy, Autism Therapy and Research Center of Excellence, Frankfurt am Main, Germany; Ernst Strüngmann Institute for Neuroscience in Cooperation with the Max Planck Society, Frankfurt, Germany Brain and Behavior Group; Dr. Senckenberg Institute of Neurooncology, University Hospital Frankfurt, Frankfurt am Main, Germany; Edinger Institute, Institute of Neurology, Goethe University, Frankfurt am Main, Germany; Frankfurt Cancer Institute, Goethe University, Frankfurt am Main, Germany; Freie Universität Berlin, AG Brain and Behavior, Berlin, Germany

**Author notes:** These authors contributed equally to this work and share first authorship (Julia Schwarzpaul, Clara M. Droell).

## Abstract

The 16p11.2 gene *QPRT*, encoding a key enzyme of the kynurenine pathway, has been linked to neurodevelopmental disorders including autism spectrum disorder (ASD). To investigate its role in early human brain development, we inhibited QPRT in stem cell-derived cerebral organoids. QPRT inhibition resulted in reduced organoid size, driven by premature neural differentiation resulting in depleted progenitor populations. Single-cell transcriptomics revealed an excitation/inhibition imbalance, with reduced excitatory and increased inhibitory neuron populations. We observed metabolic stress signatures, including pseudo-hypoxia, oxidative stress, and mitochondrial dysfunction, likely linked to NAD^+^ depletion and QUIN accumulation following QPRT inhibition. Notably, downregulation of *LHX2* and *PRDX1* may underlie impaired neural patterning and excitotoxic vulnerability. In addition, we report astrocytic and radial glia dysfunctions, indicating broad effects across multiple cell types. Disease gene enrichment analyses showed significant overlap with ASD-associated genes, especially during early differentiation. These findings suggest the loss or reduction of QPRT to shift neural development and neuronal homeostasis towards an imbalance in excitatory and inhibitory neuronal populations, a mechanism previously associated with neurodevelopmental disorders.

## Introduction

Neurodevelopmental disorders (NDDs) encompass a heterogeneous group of conditions with onset in early development, including autism spectrum disorder (ASD), intellectual disability (ID) as well as neurological conditions such as epilepsy (EP; Lord et al., 2018). These disorders are largely genetically driven and have a high comorbidity; for example, ASD is often accompanied by ADHD, epilepsy, depression, or intellectual disability (Warrier et al., 2020).

Over 1,500 genes have been implicated in NDDs, with both *de novo* and inherited, common and rare variants contributing to their etiology. Among the most frequently identified genomic variants are copy number variations (CNVs) of chr16p11.2, which are associated with neurodevelopmental as well as neurological disorders (REF). Both deletions and duplications of this region are associated with an increased liability for ASD, intellectual disability, developmental delay, schizophrenia, and bipolar disorder (Woodbury-Smith and Scherer, 2018). Specifically, deletions of 16p11.2 increase susceptibility for ASD approximately nine-fold (Moreno-De-Luca et al., 2015; Stein, 2015). The whole region has been studied in animal models, including zebrafish and rodents, giving valuable insights on the developmental and behavioral phenotypes of the associated disorders (Golzio et al., 2012; Pucilowska et al., 2015; Openshaw et al., 2023). More recently, researchers have studied 16p11.2 CVNs in induced pluripotent stem cell (iPSC)-derived neurons. In contrast to animal models, these allow studying disease phenotypes in a human genetic background. Neurons differentiated from iPSCs of 16p11.2 CNV carriers exhibited reduced synaptic density, smaller size, and shortened dendrites (Deshpande et al., 2017; Roth et al., 2020). Furthermore, cortical organoids from deletion carriers showed an increased size, a higher neuronal-to-progenitor ratio as well as altered synaptic and ion channel pathways; duplication carrier-derived cortical organoids showed a microcephaly phenotype (Urresti et al., 2021; Tai et al., 2022; Fetit et al., 2023; Kostic et al., 2023). Various cell types of the human brain including progenitor cells, inhibitory and excitatory neurons as well as glia have been described to be affected in NDDs, alongside metabolic deficits including mitochondrial dysfunction (Fetit et al., 2021; Allen et al., 2022; Russ et al., 2025). Similarly, cellular and metabolic abnormalities potentially also contribute to the altered neuronal-to-progenitor ratios and synaptic phenotypes observed in 16p11.2 deletion cortical organoids, likely driven by individual genes of the whole region.

With a total size of ∼600kb, the 16p11.2 region encompasses 29 genes – however, their individual contribution to the etiology of the various distinct symptoms is to date largely unknown. While several genes are considered as key candidates for specific phenotypes, none of them individually explains the full phenotypic spectrum. The genes *potassium* c*hannel tetramerization domain containing 13 (KCTD13), mitogen-activated protein kinase 3* (*MAPK3)* or *TAO kinase 2 (TAOK2)* have been in the focus of functional studies of the region 16p11.2. However, many genes of the region are still understudied (Calderon De Anda et al., 2012; Yufune et al., 2015; Arbogast et al., 2016; Richter et al., 2019). The 16p11.2 gene *quinolinate phosphoribosyltransferase (QPRT)* has emerged as a gene of interest, as it is strongly upregulated at the onset of neuronal differentiation of human neuroblastoma cells and is essential for brain-derived neurotrophic factor (BDNF)-mediated neuronal maturation (Haslinger et al., 2018; Neves et al., 2022). QPRT functions within the kynurenine pathway (KP), the primary route for tryptophan degradation and NAD^+^ synthesis (Braidy et al., 2011). By converting quinolinic acid (QUIN) into nicotinic acid, QPRT prevents the accumulation of QUIN, which can induce NMDA receptor-mediated excitotoxicity. Inhibition of QPRT, whether genetically or by the competitive inhibitor phthalic acid (PA), reduces NAD^+^ levels, impairs neurite outgrowth, and compromises neuronal survival (Haslinger et al., 2018; Neves et al., 2022). Excess QUIN has also been implicated in several neurodegenerative and inflammatory conditions, including Alzheimer’s disease, Huntington’s disease and multiple sclerosis (Braidy et al., 2011; Savitz, 2020).

Here, we aimed at elucidating how QPRT is shaping human neurodevelopmental trajectories and their potential involvement in 16p11.2-associated disorders, specifically ASD. We employed stem cell-derived cerebral organoids (COs), as they intrinsically form a high variety of neural cell types, allowing to identify vulnerable cell populations during the trajectory of CO development (Lancaster et al., 2013). We inhibited the encoded QPRT protein using 5 mM phthalic acid throughout differentiation and conducted longitudinal analyses of CO development, integrating gross morphological and immunocytochemical assessments with transcriptomic profiling via bulk RNA sequencing at multiple stages and single-cell RNA sequencing of mature organoids.

## Material and Methods

### Stem cell maintenance

H9 wild-type (WT; WA09, WiCell) human embryonic stem cells were maintained on plates coated with Corning® Matrigel® hESC-qualified Matrix (354277) in Essential 8 Flex (Gibco) medium at 37 °C with 5% CO2 in a humidified incubator.

### Generation of cerebral organoids and PA treatment

Cerebral organoids were generated following Esk et al., 2020, with slight modifications (Esk et al., 2020); also see Supplementary Methods). Per well of a U-bottom 96-well plate (Biofloat, Sarstedt), 2,500 cells were seeded in E8 Flex medium. Two batches of H9 WT cerebral organoids were generated following the same procedure (Batch A and B). 5 mM Phthalic acid (PA; Sigma) was added starting on day 3 to chemically inhibit QPRT. PA was dissolved directly in the respective media (E8 Flex, NIM, IDM-A, or IDM+A) to generate the 5 mM PA media, and the pH was adjusted to match the control medium (0 mM PA).

### Organoid dissociation for bulk RNA sequencing

For bulk RNA sequencing, two COs per batch, condition and time point were collected. COs were dissociated using 20 U/ml papain (Merck) and 125 U/ml DNase I (Merck; also see Supplementary Methods). Organoids were minced with pipette tips to aid digestion, incubated for 30 minutes at 37 °C on a shaker at 150 rpm and triturated afterwards to reduce clumping. Additional incubation and trituration were applied, if necessary. Dissociated cells were pelleted and stored at - 80 °C until RNA extraction.

### Single-cell RNA sequencing

On day 112, one organoid from the 0 mM PA condition and one from the 5 mM PA condition were collected. Whole, viable organoids were transferred into 5 ml tubes with their respective media, kept on ice, and shipped to Singleron Biotechnologies (Cologne) overnight for dissociation, library preparation, and sequencing.

### Tissue dissociation and single-cell isolation for single-cell RNA sequencing

At Singleron, ice-cold organoids were centrifuged at 350×g for 2 minutes, washed with 5 ml PBS, and centrifuged again at 350×g for 2 minutes. COs were resuspended in 800 µl of a 1:1 dilution of sCelLiVE® Tissue Dissociation Solution in PBS, followed by trituration for 30 seconds using a P200 pipette. The suspension was incubated at 37 °C at 500 rpm for 5 minutes, with additional trituration every 30 seconds to achieve complete dissociation. The process was monitored under a light microscope. After dissociation, the suspension was filtered through a 40 µm strainer, centrifuged at 350×g for 5 minutes at 4 °C. The resulting single-cell pellet was resuspended in 1 ml cold PBS. Cell counts were performed using propidium iodide and a Luna FX7 automated cell counter (Logos Biosystems, Villeneuve d’Ascq, France).

### Single-cell RNA sequencing library preparation

A total of 22,500 cells were loaded onto a microfluidic chip using the Singleron Matrix® Single Cell Processing System (Singleron Biotechnologies), aiming to capture approximately 3,000 cells. In brief, barcoded beads carrying unique cell barcodes were loaded onto the chip, and cells were lysed. Polyadenylated RNA from lysed cells was captured on the beads via poly(dT) sequences. The beads with bound mRNA were then collected for reverse transcription to generate cDNA, which was subsequently amplified and quality checked. Illumina-compatible sequencing libraries were prepared and sequenced on an Illumina NovaSeq 6000 platform with paired-end 150 bp reads. Sequencing reads were demultiplexed on Illumina’s BaseCloud, and the resulting fastq files were used for downstream data analysis.

### Single-cell RNA sequencing data processing and quality control

Raw Illumina fastq files were processed into gene expression matrices using CeleScope™ version 1.14.0 (www.github.com/singleron-RD/CeleScope; Singleron Biotechnologies). Reads were demultiplexed based on cell barcodes and unique molecular identifiers (UMIs). Adapter sequences and poly-A tails were removed using cutadapt (https://cutadapt.readthedocs.io/en/stable/installation.html), and trimmed Read2 sequences were aligned to the GRCh38 human genome with Ensembl version 92 gene annotations using STAR v2.6.1a_08-27 and featureCounts 2.0.1. Reads sharing the same cell barcode, UMI, and gene were combined to calculate UMI counts per gene per cell. The resulting UMI count tables for each cell barcode were then used for further analysis.

### Single-cell RNA sequencing analysis

Cell-by-gene count tables were analyzed in R (version 4.3.1) using standard pipelines (Seurat 5.0.3, DoubletFinder 2.0.3, DropletUtils 1.20, harmony 1.2.0, with the respective dependencies). Genes detected in less than three cells and cells expressing less than 250 genes were removed. Low-quality cells (mitochondrial RNA >20%, ribosomal RNA <5%) and doublets were excluded (Supplementary Figure 1). The remaining data were normalized, scaled, and highly variable features identified. Harmony was used for batch integration (SCTransform based integration), and harmony and elbow-plotting determined meaningful components (n=30) for clustering (Seurat built in functions FindNeighbors FindClusters). Clusters were optimized based on resolution and results visualized using Clustree. Stable clusters were identified at res=0.6. Similarly, UMAP parameters were optimized for cluster visualization (min distance 0.3, spread 4). Cell cycle states were scored using the built in Seurat function (Supplementary Figure 2). Cluster marker genes (Seurat FindAllMarkers) were identified (log2FC>0.25). Cluster identity was confirmed by mapping to a reference cerebral organoid dataset (Li et al., 2023; Supplementary Figure 3) using Scater (v 1.28).

Differentially expressed genes (DEGs) across conditions were identified for each Cluster with multiple testing correction for all genes tested. Functional enrichment analysis was performed to identify associated biological processes using gprofiler2 (v 0.2.3).

### RNA extraction for bulk sequencing

For bulk gene expression analysis, RNA was isolated using the NucleoSpin RNA kit (Macherey-Nagel) according to the manufacturer’s protocol. The RNA was eluted in RNase-free water, quantified with a Nanophotometer (Implen), and sent to Life and Brain GmbH (Bonn) for bulk sequencing using the QuantSeq 3′-mRNA Library on a NovaSeq 6000 with 10Mio Raw Reads.

### Analysis of bulk sequencing

Raw FastQ reads were quality-checked and trimmed using Trimmomatic with default settings. Reads were aligned to the hg38 genome with Entrez annotations using the align() function from Rsubread. Count matrices were generated via featureCounts(), and redundant annotations were consolidated by summing counts. Genes with no variance or detected in fewer than 25% of samples were excluded, leaving 13,185 genes for downstream analysis and an average read count per sample of 5,7 Mio.

Replicate consistency and identification of technical outliers was assessed through hierarchical clustering (Ward.2), Euclidean distances, PCA, and multidimensional scaling, confirming biological replicates and clear clustering by differentiation stage. For differential expression analysis, raw counts were imported into DESeq2 using DESeqDataSetFromMatrix(), quantile-normalized, and log2(count+1) transformed. Two models were applied: Model A: read ∼ 1 + as.factor(PA_treatment) + (1|Batch) for individual time points, and Model B: read ∼ 1 + Day*as.factor(PA_treatment) + (1|Batch) for combined effects. Batch effects were modeled as random factors. Differential expression was analyzed with lmFit() and statistical significance estimated via empirical Bayes moderation (eBayes()), with FDR < 0.05 considered significant.

Gene set enrichment analysis of differentially expressed genes (DEGs) was performed using gprofiler2 with the gSCS correction to account for GO term hierarchy. Only DEGs with FDR < 0.05 were included.

Cell type composition in bulk RNA-seq samples was estimated using deconvolution (MuSiC v 1.0 (Fan et al., 2022) with single-cell RNA-seq data from the day-112 cerebral organoids (9,107 cells, 21,419 genes) as well as cells on published cerebral organoid data (Li et al., 2023). Proportions were inferred using the music_prop() function.

### Disorder gene list enrichment testing

Differentially expressed genes from single-cell and bulk RNA sequencing were tested for enrichment among curated candidate gene lists for autism spectrum disorder (ASD), epilepsy (EP), and intellectual disability (ID). Specifically, we included ASD-associated genes curated by the SFARI database (August 19, 2024 release), ASD-associated co-expression modules M12 (neuronal) and M16 (astrocyte/microglia) derived from post-mortem brain tissue (Voineagu et al., 2011), rare de novo variant (RDNV) gene sets (De Rubeis et al., 2014; Iossifov et al., 2014), curated monogenic epilepsy genes (Oliver et al., 2023; EpilepsyGenes_v2024_03), and ID risk genes (Liu et al., 2018). The complete lists are provided in Supplementary Table 1.

### Morphological analysis and immunostaining

Bright-field imaging and immunostaining of cerebral organoids were performed to assess morphological features, cell type composition, and neuronal subtypes. Embryoid bodies and organoids were imaged using a Motic AE31 microscope to monitor potential size differences. For histological analysis, three organoids per time point, batch, and condition were fixed in 4% PFA for 3 h at room temperature, washed in PBS, and stored in 30% sucrose prior to embedding in Tissue-Tek O.C.T. COs were cryosectioned at 20 µm using a Leica CM3050 S cryostat and mounted on Superfrost Plus slides. Antigen retrieval was performed in citrate buffer (pH 6.0) at 95 °C for 20 min, followed by permeabilization and blocking in 0.05% Triton X-100 with 4% normal goat serum for 30 min. Primary antibodies were applied overnight at 4 °C, followed by 1 h incubation with secondary antibodies at room temperature, and slides were mounted in Fluorescent Mounting Medium (Dako). Two antibody panels were used: one for assessing neural progenitors (Paried Box Protein 6 (PAX6); Rabbit; Proteintech, 12323-1-AP; 1:500), mature neurons (Microtubule-Associated Protein 2 (MAP2); Mouse; Millipore; MAB3418; 1:500), glial cells (Glial Fibrillary Acidic Protein (GFAP); Guinea pig; Synaptic systems; 173 004; 1:500), and general morphology with Hoechst 33342 staining (Thermo Fisher); and a second panel for distinguishing glutamatergic (vesicular Glutamate Transporter 1 and 2 (vGlut1/2); Mouse; Sigma; AMAb91041 (vGlut1); AMAb91086 (vGlut2); 1:500) and GABAergic neurons (Glutamate Decarboxylase 65 and 67(GAD65/67); Rabbit; Sigma; G5163) in combination with MAP2 (Guinea Pig; Synaptic systems, 188 044) and Hoechst 33342. For each condition, slices from two organoid batches across days 25, 40, 60, and 80, treated with 0 or 5 mM PA, were included as biological replicates, with technical duplicates per staining. Slides were imaged using an Axioscan 7 (Zeiss) with ZEN blue software (v3.7), scanning all slides from a given time point in a single session under consistent settings.

### Image Analysis

Images of cerebral organoid slices that were severely overlapped, damaged, or misaligned on the slides were excluded from analysis. Remaining images were blinded using a custom R script and analyzed in Zeiss ZEN software (v3.9.3) using the “Draw Contour (Spline)” tool. Contours were manually drawn around each organoid slice and any large holes or cracks were subtracted to calculate the accurate area. For each slice, we quantified the total area (µm^2^) and measured the mean fluorescence intensity of MAP2, PAX6, GFAP, and Hoechst 33342. Neural rosettes were similarly outlined, focusing only on clearly defined regions, with the same set of parameters recorded.

For glutamatergic and GABAergic neurons (vGlut1/2, GAD65/67, MAP2), images were analyzed automatically in ImageJ (Fiji v1.54i) using a custom macro (see Supplementary Methods). All measurements were compiled in Excel, unblinded, and statistical comparisons between 0 and 5 mM PA groups were performed for each time point using Wilcoxon rank-sum tests with Bonferroni correction in R. Significant results were further assessed for batch effects using a linear mixed-effects model.

### Data availability

Sequencing raw data are available on request.

### Ethics statement

The use of human embryonic stem cells in this study was authorized by the Robert Koch Institute, Germany (3.04.02/0174).

## Results

### QPRT inhibition impairs organoid growth

QPRT inhibition via 5 mM PA (Figure 1A) led to a significant decrease in organoid size (Figure 1B, C). While sizes of treated and untreated embryoid bodies were mostly comparable within batches prior to neural induction, we observed profound size differences during early differentiation in the QPRT-inhibited COs after Matrigel-embedding and CHIR 99021 treatment (p < 0.05 for both batches for d16, d17, d20). The differences in differentiation and CO development were also notable in organoid morphology. The organoids treated with PA descriptively showed a reduced neural rosette formation, suggesting neural progenitor depletion, and an increase of outgrowing cells from around d25 on (Figure 1D).

**Figure 1:**
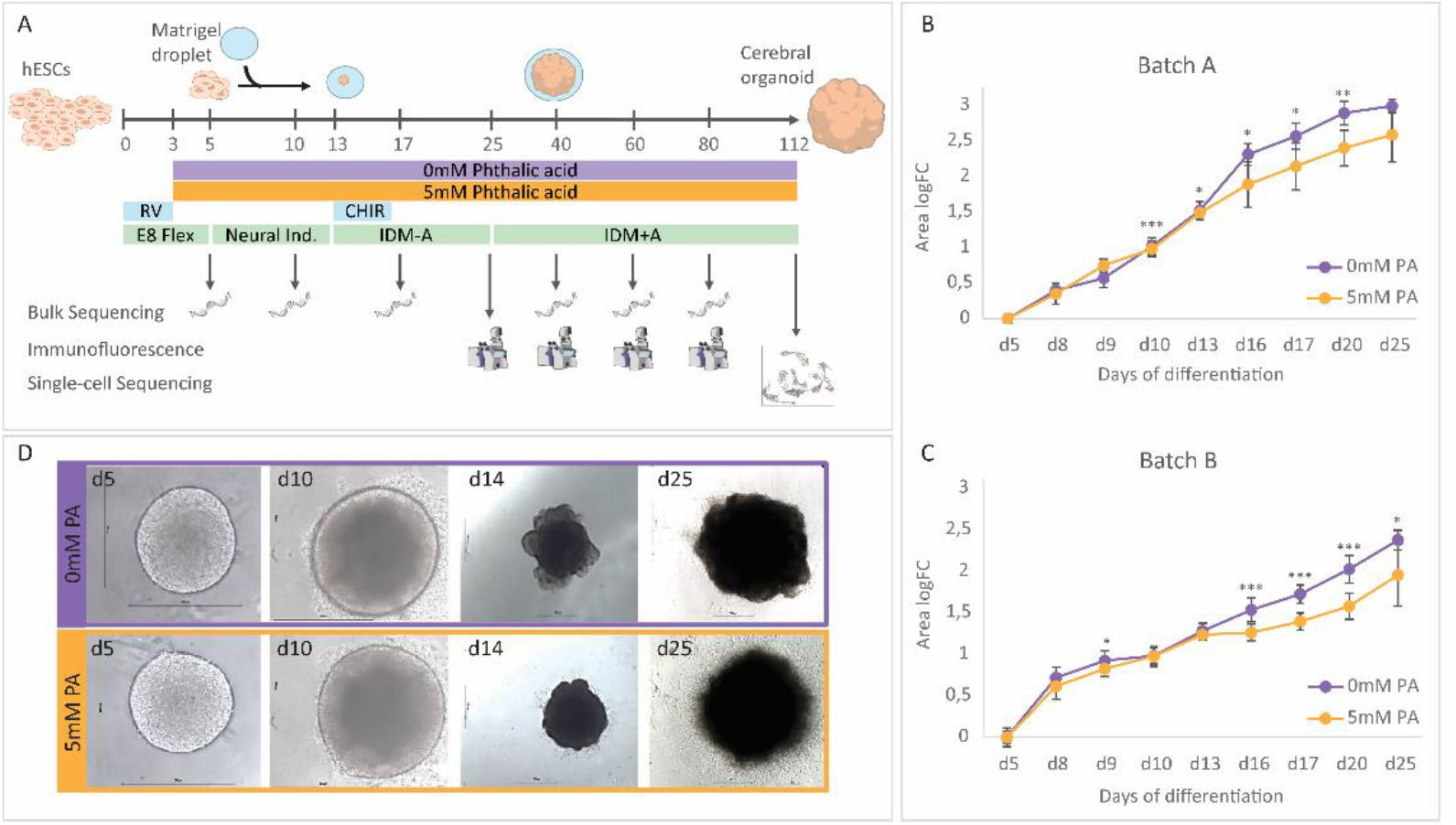
**A)** Overall study design: Cerebral organoids were differentiated following a protocol modified from Esk et al. 2020 (Esk et al., 2020, Lancaster et al., 2013). COs were treated with 0 (control) or 5 mM phthalic acid (QPRT-inhibited) starting from d3 on. **B)** and **C)** Administration of 5 mM PA impaired organoid growth after neural induction, replicated across batches. **D)** QPRT inhibition altered organoid morphology and impaired organoid integrity by outgrowing cells. RV: RevitaCell (Gibco), IDM-A: Improved NeuroDMEM-A, IDM+A: Improved NeuroDMEM+A, PA: phthalic acid. * p<0.05, ** p<0.01, *** p<0.001.

### QPRT inhibition leads to differential gene expression across all cell types

To identify pathway changes induced by the inhibition of QPRT, we performed scRNA sequencing of mature organoids at d112. Of 21,419 genes analyzed after quality control, 1,034 were differentially expressed across all cells/in any cluster when comparing the 0 mM and 5 mM PA condition for all cell types together (271 downregulated, 763 upregulated, FDR < 0.05, Supplementary Table 2). Genes upregulated in the treated condition were associated with GO terms related to synaptic signaling (p < 0.05, Supplementary Figure 4; Supplementary Table 3). Multiple differentially expressed genes (DEGs) encoded GABA receptor subunits such as the ASD-associated genes g*amma-aminobutyric acid type a receptor subunit gamma 3* (*GABRG3;* log2FC = 1.055, FDR = 0.032) and g*amma-aminobutyric acid type a receptor subunit beta 2* (*GABRB2;* log2FC = 0.51, FDR = 0.002). The gene e*rb-b2 receptor tyrosine kinase 4* (*ERBB4*), expressed by GABAergic neurons (Asede et al., 2021), was upregulated (log2FC = 1.17, FDR = 5.91E-61). The strongest upregulation was seen for *endothelin 1* (*EDN1*; log2FC = 4.64, FDR = 1.00E-12), which is expressed by neurons as well as astrocytes under stress conditions (Hasselblatt et al., 2001). Further, the set of upregulated genes showed strong enrichment for ASD candidate genes (07_SFARI_all: OR = 1.98, FDR = 1.06E-05; 08_ASD_M12_Voineagu_2011: OR = 2.12, FDR = 6.90E-03; 11_ASD_RDNV_DeRubeis_2014: OR = 3.56, FDR = 0.03; Supplementary Table 4).

Genes downregulated upon QPRT inhibition were enriched for processes connected to brain development, such as cerebral cortex development (p = 3.59E-06), including the EP-associated gene *neuronal differentiation 2* (*NEUROD2;* log2FC = -0.595, FDR = 1.761E-26) and the ASD- as well as ID-associated gene *forkhead box P2* (*FOXP2;* log2FC = -1.574, FDR = 0.009). Additionally, the GO term neural precursor cell proliferation (p = 2.66E-05) with genes like *paired box 6* (*PAX6;* log2FC = -0.548; FDR = 4.29E-04; implicated in ASD, EP and ID) was associated with downregulated DEGs. In contrast to the upregulated genes, downregulated ones were only associated with an ASD-implicated gene set involved in astrocytes and microglia regulation (09_ASD_M16_Voineagu_2011: OR = 4.27, FDR = 4.48E-06, Voineagu et al., 2011). While we do see overlaps with nominal significance for EP- and ID-associated genes, none of those reached significance after correction for multiple testing. Among the top downregulated ones upon PA treatment were the ASD genes *peroxiredoxin 1*(*PRDX1;* log2FC = -0.67, FDR = 2.40E-60) and *LIM homeobox* 2 (*LHX2;* log2FC = -1.00, FDR = 1.61E-54), the latter is crucial for nervous system development and associated with variable neurodevelopmental phenotypes (Schmid et al., 2023). Further, we found *fatty acid binding protein 7* (*FABP7*; log2FC = -0.95, FDR = 3.56E-120) to be downregulated, a gene with an important role in establishing the radial glia fiber system as well as cortical layer formation, and recently discussed in the context of ASD etiology (Han et al., 2025). Taken together, upregulated genes upon QPRT inhibition are associated with synaptic signaling and overlap with ASD candidate genes, including neuronal-associated genes. For downregulated genes we saw an association with nervous system development and limited overlap with ASD-associated markers for astrocytes and activated microglia. Finally, the downregulation of genes associated with neural cell precursor proliferation relates to the reduced organoid size and confirms the hypothesis of progenitor depletion.

### In mature organoids, QPRT inhibition leads to altered cell type distribution and distinct changes in gene expression patterns

We then investigated altered cell type distributions and pathway changes within individual cell types. We identified 14 distinct cell types in 112d old organoids (Clusters 0-13; Figure 2A, Supplementary Figure 3). QPRT inhibition did not give rise to any different cell type. At this mature stage, we detected radial glia and outer radial glia (RG_oRG; Cluster 3 and 10) as well as cycling radial glia (ccRG; Cluster 4 and 9), indicating an ongoing progenitor proliferation and potential differentiation of the organoids at later stages. Additionally, we identified astrocytes (Ast; Cluster 6), which are expected to arise only after 60 to 80 days of differentiation (Porciúncula et al., 2021). Neural progenitors were found for excitatory (Ex_P_Cl5) and inhibitory (In_P_Cl8) cells. We further detected excitatory neurons (Ex_N_Cl 1, 2, 7 and 11) and inhibitory neurons (In_N_Cl 0, 12 and 13) with a 70:30 ratio in the reference organoid, which resembles the natural state of the fetal brain (Porciúncula et al., 2021). Within the excitatory neuron clusters, we identified neurons from the cortical layers VI up to layer II which emerges last during cortical development (Vanderhaeghen and Polleux, 2023).

**Figure 2:**
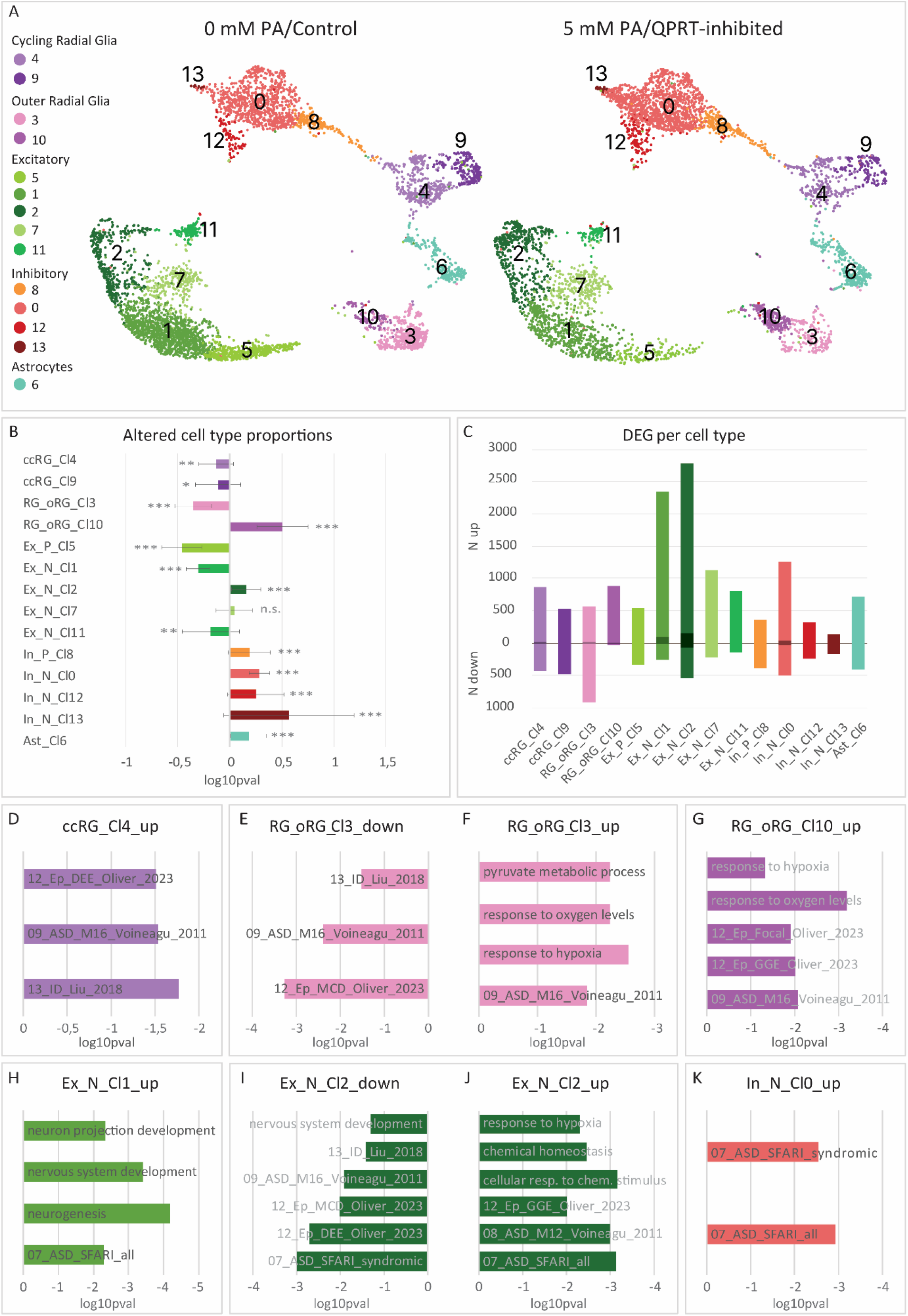
**A)** and **B)** QPRT inhibition significantly altered cell type frequencies in mature organoids. **C)** Numbers of differentially expressed genes in distinct cell types. Lighter shade: p < 0.05, darker shade: FDR < 0.05. **D)-K)** Enrichment of cell type-specific DEGs with GO terms or disease gene lists. All p < 0.05. ccRG: cycling radial glia; RG_oRG: radial glia/outer radial glia; Ex_P: excitatory progenitors; Ex_N: excitatory neurons; In_P: inhibitory/intermediate progenitors; In_N: inhibitory neurons; Ast: astrocytes DEG: differentially expressed genes. ns: not significant, * p<0.05, ** p<0.01, *** p<0.001.

QPRT inhibition significantly altered cell type distribution (Figure 2A, Supplementary Table 5). While PA treatment led to a decreased proportion of both cycling radial glia clusters (ccRG_Cl4/9), the proportion of outer radial glia (oRG_Cl3/10) shifted: Cluster 3 decreased while cluster 10 increased. Further, QPRT inhibition led to a significant increase of astrocytes (Ast_Cl6). Interestingly, inhibitory neuron clusters showed an increased proportion in the treated condition while excitatory neuron clusters decreased. The inhibitory cluster 13 showed the biggest change in proportion (OR = 3.69, FDR = 9.02E-05). Combined with the significant increase in the remaining inhibitory clusters 0, 12 as well as their progenitor cluster 8, this confirms our observation of an enrichment of GABA-associated genes in the DEG analysis across all cell types. For the excitatory progenitor cluster 5 we observed the strongest decrease. Additionally, the excitatory clusters 1 and 11 were significantly reduced upon QPRT inhibition. Taken together, QPRT inhibition led to an altered ratio of excitatory to inhibitory neurons: While there was a ∼70:30 ratio of excitatory (Clusters 1, 2, 7, 11) to inhibitory (Clusters 0, 12, 13) neurons in the untreated organoid (n_Ex_N_ = 1967; n_In_N_ = 928), the ratio dropped to 50:50 in the QPRT-inhibited condition (n_Ex_N_ = 1370; n_In_N_ = 1358).

When testing for DEGs across distinct cell types (Figure 2C, Supplementary Table S6) we observed an excess of overexpressed genes in excitatory neurons of Cluster 1 and 2 (Cluster 1: N_p_ = 2252, N_FDR_ = 90; Cluster 2: N_p_ = 2627, N_FDR_ = 150). In cycling radial glia (ccRG_Cl4, Figure 2D) we found an enrichment of candidate genes for ASD, EP as well as ID in genes upregulated upon QPRT inhibition (Figure 2D, Supplementary Tables 7 and 8). However, specific biological processes could not be associated. Interestingly, the downregulated gene set of outer radial glia showed a similar pattern (oRG_RG_Cl3, Figure 2E). Genes upregulated within this cell type were associated with the response to hypoxia (Figure 2F). Similarly, we observed an enrichment of hypoxia-related genes for the upregulated gene set of the other radial glia cluster (RG_oRG_Cl10, Figure 2G). For the excitatory neuron clusters 1 and 2 which had the highest number of DEGs, we again report an overlap with hypoxia-related genes in the upregulated genes of Ex_N_Cl2 (Figure 2H, I, J), while upregulated genes of Ex_N_Cl1 showed a strong association with neurogenesis-related GO terms. In the case of inhibitory neurons, upregulated genes of Cl0 showed a strong enrichment of genes associated with ASD (Figure 2K). Genes differentially regulated in astrocytes or inhibitory/excitatory neuronal progenitors showed no significant overlap with any of the disease gene lists. While the list of DEGs in astrocytes was not enriched for a specific GO term, we did see an association with the biological process of glycolysis and gluconeogenesis via the significant downregulation of the central glycolysis gene *GAPDH* specifically within this cell type (Supplementary Tables 6 and 7).

### QPRT inhibition impacts on distinct cell type development throughout neural development

In addition, we analyzed transcriptional changes over time, which, in combination with the single-cell sequencing analysis of mature COs, allowed us to evaluate the dynamic cell type compositions.

Overall, cell type compositions changed over time as expected, i.e., progenitor cell types decreased towards later differentiation, while cells of neural identity increased (Figure 3A). Of the 14 identified cell types, 6 showed significant PA-induced changes at one or more timepoints.

**Figure 3:**
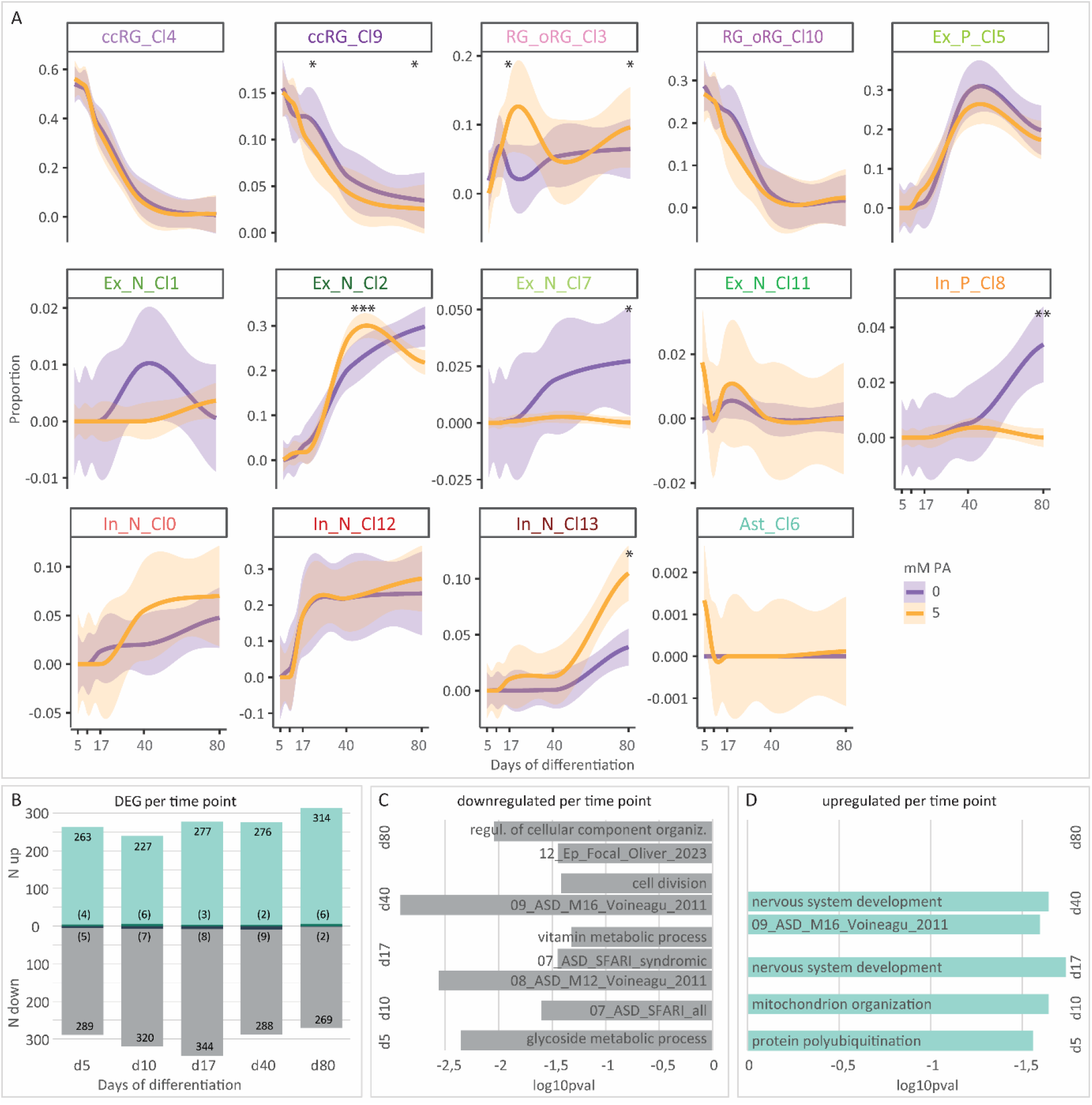
**A)** QPRT inhibition affects cell type development over time. **B)** Number of genes differentially regulated upon QPRT inhibition per time point. Lighter shade: p < 0.05, darker shade: FDR < 0.05. **C)** and **D)** GO terms and disease gene lists enriched for differentially regulated genes per time point. All p < 0.05. ccRG: cycling radial glia; RG_oRG: radial glia/outer radial glia; Ex_P: excitatory progenitors; Ex_N: excitatory neurons; In_P: inhibitory/intermediate progenitors; In_N: inhibitory neurons; Ast: astrocytes; PA: phthalic acid; DEG: differentially expressed genes. * p<0.05, ** p<0.01, *** p<0.001.

In both conditions, cycling radial glia (ccRG_Cl9) showed a strong decrease over time. However, in the treated condition, we observed a transient plateau of ccRG_Cl9 proportion, resulting in a significant increase at d17 (p = 1.61E-02) as well as on d80 (p = 4.67E-02). Outer radial glia (RG_oRG_Cl3) of the treated condition showed a strong increase in proportion during early differentiation (d17, p = 2.13E-02), followed by a slight drop mid maturation at d40 (p > 0.05) and again a significant increase towards late maturation at d80 (p = 4.39E-02), confirming the findings of the d112 scRNA sequencing. Notably, two clusters corresponding to excitatory neurons (Ex_N_Cl2 and Ex_N_Cl7) displayed distinct temporal dynamics. For cluster 2 we found a significant overshoot in the QPRT-inhibited organoids at d40 (p = 1.36E-04), followed by a pronounced decrease towards later maturation. The proportion of excitatory neurons of cluster 7 in the untreated group showed a continuous increase starting at d17, however, upon QPRT inhibition this cell type remained at low levels over time resulting in a significant decrease at d80 (p = 4.63E-02). This again validates our scRNA finding of a reduction of excitatory neurons at d112.

Regarding inhibitory neurons, in QPRT-inhibited cells we observed a plateau of intermediate progenitors (In_P_Cl8) while they increased over time in the control organoids, resulting in a significant difference at d80 (p = 2.80E-03). Finally, for inhibitory neurons (In_N_Cl13) we observed a significantly increased proportion in the treated condition at day 80 (p = 2.16E-02), also reflecting our scRNA findings at d112. In summary, the overall proportions of different neural cell types and the finding of a decrease of excitatory and an increase of inhibitory neurons upon inhibition of QPRT was confirmed in both independent analyses. The reduced proportion of cycling radial glia (ccRG_Cl9) together with the transient overshoot of the excitatory neuron subset (Ex_N_Cl2) further point towards a premature neural differentiation during QPRT inhibition.

### QPRT inhibition alters neuronal development and accelerates differentiation

We then looked at differentially expressed genes for the investigated time points (Figure 3B, Supplementary Table 9). We observed a similar amount of DEGs for all the analyzed time points, with the top number of downregulated genes at d17 and the top number of upregulated ones at d80.

On day 5, prior to neural induction, 289 genes were downregulated (Figure 3B), showing enrichment for the GO term glycoside metabolic process (Figure 3C, Supplementary Table 10), suggesting an early onset of response to hypoxia stress. Among the downregulated genes, five exhibited an FDR < 0.05 (Figure 3C). Notably *cerebellin 1 precursor* (*CBLN1*; logFC = -5.82, FDR = 0.031), which is critical for maintaining synaptic integrity and plasticity (Matsuda and Yuzaki, 2011), and *rho GTPase activating protein 39* (*ARHGAP39*; logFC = -5.34, FDR = 0.042), which plays a role in postsynaptic organization at glutamatergic synapses (Ito and Nagata, 2022). In contrast, a total of 263 genes were significantly upregulated (p < 0.05), which were enriched for protein polyubiquitination (Figure 3D, Supplementary Table 10), indicating the targeting of proteins for degradation, potentially as early stress response. Four of the upregulated genes reached significance after correction (FDR < 0.05), unexpectedly for this early time point including *protocadherin gamma subfamily B6* (*PCDHGB6*; logFC = 5.69, FDR = 2.70E-04), a gene involved in establishing specific neuronal connections (Stelzer et al., 2016).

On day 10, i.e., during the neural induction state, 320 genes were downregulated. These did not significantly overlap with a specific GO term (Figure 3C) but were enriched for ASD-associated genes (07_ASD_SFARI_all: OR = 1.71, p = 0.03). Of the seven genes showing an FDR < 0.05 (Figure 3B), three were involved in cytoskeleton regulation: *Rotatin* (*RTTN;* logFC = - 5.98, FDR = 0.04; Chou and Tang, 2021), *LEM domain containing 3* (*LEMD3;* logFC =-5.90, FDR = 0.049; Chambers et al., 2024) and the ASD-, EP- and ID-associated gene *spectrin repeat containing nuclear envelope protein 1* (*SYNE1*; logFC = -5.59, FDR = 0.01). In contrast, 227 genes were upregulated (Figure 3B), with an association with mitochondrion organization (Figure 3D), suggesting a response to cellular stress.

On day 17, the time right after starting neuroectoderm expansion, 344 genes were downregulated and enriched for vitamin metabolic processes (Figure 3C). Eight of these downregulated genes showed an FDR < 0.05, including the NDD-associated gene *catechol-o-methyltransferase (COMT)*, which is involved in the methylation of dopamine (logFC = -5.84, FDR = 7.82E-04; Papaleo et al., 2016), and *serine/threonine kinase 32C* (*STK32C*), a gene highly expressed in the brain (logFC = -4.62, FDR = 0.046; (Dempster et al., 2014). For the genes downregulated upon QPRT inhibition at d17, we see an overlap with ASD-associated genes (08_ASD_M12_Voineagu_2011: OR = 2.60, p = 2.75E-03; 07_ASD_SFARI_syndromic: OR = 0, p = 0.04). Further, 277 genes were upregulated (Figure 3B), showing an enrichment for the GO term nervous system development (Figure 3D). Genes with an FDR < 0.05 include the ASD- and ID-associated gene *transforming growth factor beta receptor 2* (*TGFBR2*; logFC = 4.97, FDR = 7.82E-04).

On day 40, which is mid maturation, 288 genes were downregulated and enriched for cell division processes (Figure 3C). These genes were further enriched for ASD-associated genes (09_ASD_M16_Voineagu_2011: OR = 2.88, p = 1.20E-03). Nine genes reached an FDR < 0.05, including *leucine rich repeats and immunoglobulin like domains 2* (*LRIG2;* logFC = -5.41, FDR = 0.044), a gene known to promote epidermal growth factor signaling and enhance cellular proliferation (Hu et al., 2022). Further, 276 genes were upregulated (Figure 3B), and these genes were enriched for the GO term nervous system development (Figure 3D). Similarly to the downregulated ones, genes upregulated at d40 showed an enrichment of ASD-associated genes (09_ASD_M16_Voineagu_2011: OR = 0, p = 0.025). Among the two genes with an FDR < 0.05, *tubulin epsilon and delta complex 2* (*TEDC2*; logFC = 6.42, FDR = 0.01) was the one with the highest upregulation. This gene might be involved in neural development via binding to tubulin epsilon and delta complex 1 (TEDC1, Miyake et al., 2025).

At the mature state (d80), 269 genes were downregulated, showing an association with regulation of cellular component organization (Figure 3C). Among the nominally downregulated genes was *neuronal differentiation 4* (*NeuroD4)*, a transcription factor critical for neuronal development shown to reprogram astrocytes into functional neurons (logFC = - 4.61, p = 6.28E-05; (Wang et al., 2023). The downregulated gene set at d80 was enriched for EP-associated genes (12_Ep_Focal_Oliver_2023: OR = 7.42, p = 0.04). However, no association with ASD genes was identified. Further, a total of 314 genes were upregulated (6 with FDR<0.05; Figure 3B), which could not be associated with any specific biological processes (Figure 3D). The strongest effect was observed for *BR serine/threonine kinase 1* (*BRSK1;* logFC = 6.127, FDR = 9.12E-04), known to play a critical role in neuronal polarization (Kishi et al., 2005) and for *F-box protein 22* (*FBXO22*, logFC = 5.53, FDR = 9.12E-04), which is involved in neural development via d-serine (Dikopoltsev et al., 2014) and synaptic development and maintenance (Yang et al., 2022). In addition, the gene *tyrosine hydroxylase (TH)*, essential for dopamine synthesis, showed significant upregulation by d80 (logFC = 4.72, FDR = 0.014; Tolleson and Claassen, 2012) and was also nominally upregulated on day 40 (logFC = 5.43, p = 7.08E-04). An increase in expression was also observed for *polo like kinase 3 (PLK3)*, a gene implicated in cell cycle control (logFC = 4.59, FDR = 0.038; Xie et al., 2001).

### QPRT inhibition alters neural rosette density and leads to morphological changes throughout differentiation

As we report significant changes in gross morphology upon inhibition of QPRT and RNA analyses indicated a shift in cell distribution, we performed immunocytochemistry for further confirmation. On day 40, the number of rosettes per area was significantly lower in the QPRT-inhibited group (FDR = 3.82E-04; batch corrected p = 9.52E-05; Figure 4A). Additionally, we observed a nominally significant increase in rosettes per area at d80. Overall, rosettes per area decreased over time, as COs matured (Figure 4A, B).

**Figure 4:**
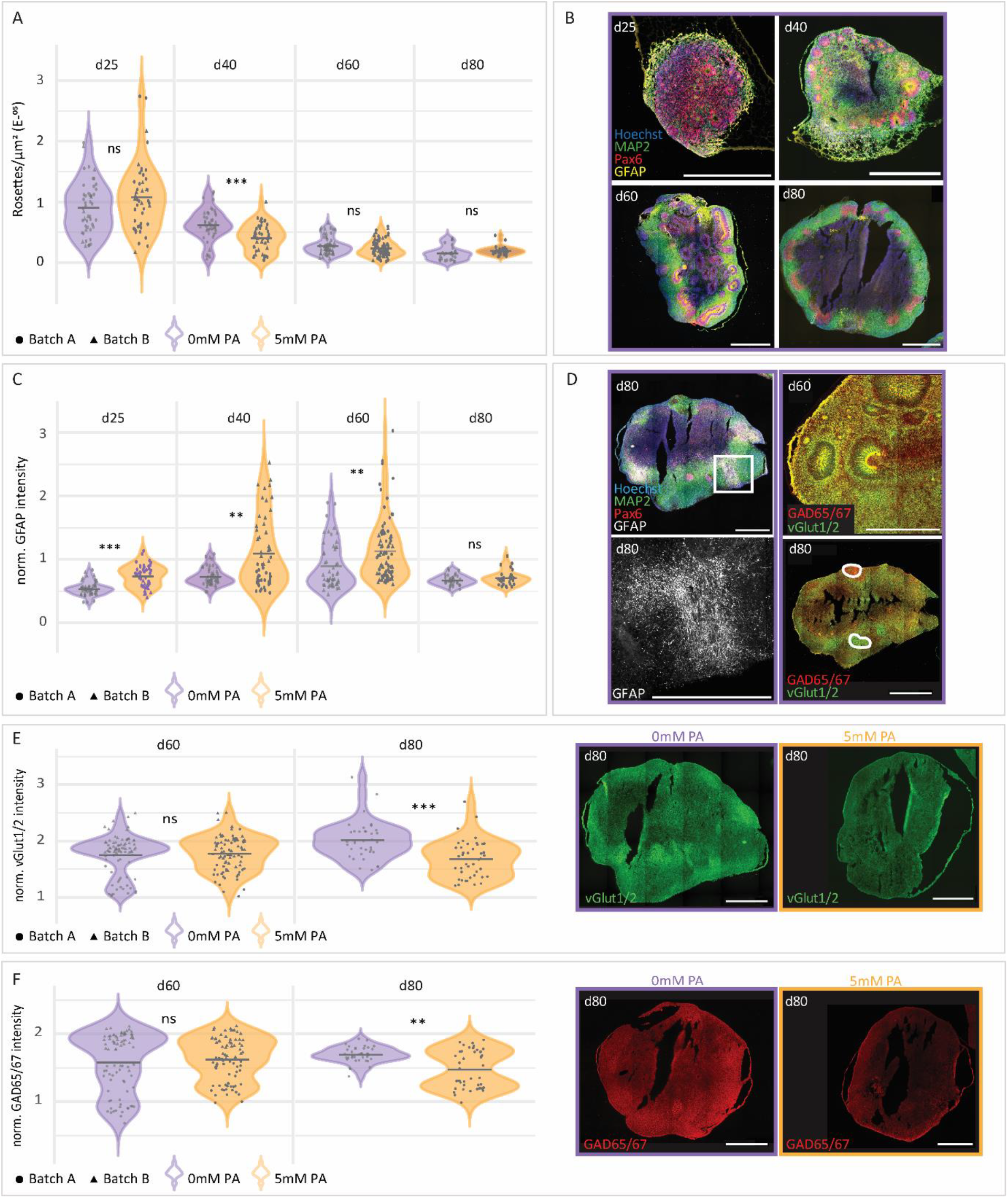
**A)** Number of rosettes per total size of organoid slice across different time points. **B)** Representative stainings of 0 mM PA CO slices over time. Scale bar: 1000 µm. **C)** GFAP intensity of organoid slices normalized to MAP2 signal at different time points. **D)** Representative stainings of 0 mM PA CO slices for GFAP as well as GAD65/67 and vGlut1/2. Scale bar: 1000 µm. **E)** Intensity of vGlut1/2 normalized to MAP2 over time, including representative stainings of 0 and 5 mM PA CO slices. Scale bar: 1000 µm. **F)** Intensity of GAD65/67 normalized to MAP2 over time, including representative stainings of 0 and 5 mM PA CO slices. Scale bar: 1000 µm. PA: phthalic acid. Ns = not significant, * = p < 0.05, ** = p < 0.01, *** = p < 0.001.

We further validated the impact on radial glia and astrocytes via GFAP staining, as observed at the transcriptomic level. Again, we report an increased GFAP intensity upon QPRT inhibition for d25 (FDR = 5.16E-08), d40 (FDR = 4.7E-03) and d60 (FDR = 0.002; Figure 4C) which remained significant after batch correction (all p <0.03). Since in cerebral organoids, astrocytes develop around day 60, the differences in earlier stages can be attributed to different numbers of GFAP^+^ radial glia cells. This again confirms the results from bulk sequencing where the proportion of outer radial glia cells was significantly higher upon QPRT inhibition at day 17 and day 80 (Figure 3A). Descriptively, and as reported in the literature (Sloan et al., 2017), GFAP^+^ cells appeared in clusters at later stages (Figure 4D).

To confirm the altered ratios of excitatory to inhibitory neurons, we additionally stained for glutamatergic markers vGlut1 and vGlut2 as well as for GABAergic markers GAD65/67. Overall, at d60 both neuronal types were identified and were mostly colocalized (Figure 4D). While the signals and statistical findings at day 60 were mixed with pronounced batch effects, findings at day 80 were in line with our transcriptomic analysis: On day 80, the signal intensity for glutamatergic markers was significantly lower in the PA-treated group (FDR = 0.004). However, no significant differences were detected for the ratio of vGlut/GAD. Descriptively, the PA-treated group exhibited lower mean ratios, indicative of a higher relative abundance of GAD65/67 compared to vGlut1 and vGlut2.

## Discussion

Here, we showed that in cerebral organoids, QPRT inhibition significantly impacts early developmental stages leading to a shift in glutamatergic/GABAergic neuron distribution, potentially attributed to an increased vulnerability to neuronal stress. At morphological level, the reduced organoid size likely was the result of the depletion of progenitor pools via premature neural differentiation, as we report genes upregulated at d17 and 40 to be associated with nervous system development (Figure 3D). In line with this, the reduction of progenitor cells was evidenced by the significant downregulation of genes associated with cell division at d40, e.g., of *LRIG2*, a gene known to promote cellular proliferation. This finding was further confirmed by a decreased number of neural rosettes at d40. At the same time, we observed a significant reduction in intermediate progenitors, whereas the mature inhibitory neurons had increased proportions, again pointing towards an accelerated differentiation. We further observed a significant downregulation of the gene *LHX2* across all cells in mature organoids at d112. Missense or gene-disrupting variants of this gene have recently been associated with ASD and microcephaly (Schmid et al., 2023). *LHX2* regulates early neural differentiation by promoting the expression of PAX6, a key neurogenic transcription factor, and by attenuating BMP and WNT signaling, thereby inhibiting non-neural fates in human embryonic stem cells (Hou et al., 2013). In derived embryoid bodies, *LHX2* was expressed in early neural rosettes, and its knockdown severely impaired their formation (Hou et al., 2013). Thus, the downregulation of *LHX2* observed here might be the causal link to the reduced neural rosette growth in the PA-treated COs, subsequently, resulting in the reduced organoid size.

QPRT inhibition significantly impacted development of several cell types, including subtypes of radial glia, excitatory and inhibitory neurons as well as astrocytes. Specifically, the reduction of excitatory and increase in inhibitory clusters were most prominent and replicated across all analyses. QPRT is catabolizing QUIN, a potent excitotoxin which binds to NMDA receptors. Hence, when QPRT is inhibited, QUIN is likely accumulating, potentially contributing to the observed reduction of excitatory neurons. The ratio of excitatory neurons overall was reduced upon QPRT inhibition, alongside an increased proportion of inhibitory neurons leading to an imbalance of excitatory to inhibitory neurons as observed in previous studies of other ASD candidate genes: Chalkiadaki and colleagues report an increased number of inhibitory neurons in parallel with a reduction of excitatory neurons in *CNTNAP2*-KO cerebral organoids (Chalkiadaki et al., 2024). Similarly, Mariani et al. observed an increase in inhibitory neurons in cortical organoids derived from iPSCs of autistic individuals (Mariani et al., 2015). These and our findings thus emphasize that altered GABAergic inhibition might be an overarching pathological mechanism. Immunostainings here confirmed the reduction in glutamatergic neurons at day 80. In contrast to the bulk sequencing results, morphological data also revealed a decrease in GAD65/67 expression. However, the markers used only identified a subset of inhibitory neurons, while the transcriptomic analyses covered the combination of various markers. Overall, these findings indicate that QPRT inhibition may lead to a dysregulation of excitatory and inhibitory neuronal ratios. Additional analyses to identify subpopulation-specific effects are needed.

Our findings repeatedly pointed towards a potential hypoxia response in the QPRT-inhibited organoids. Given that QPRT is part of the kynurenine pathway which results in NAD^+^ production, we hypothesize a depletion of NAD^+^ following QPRT inhibition leads to pseudo-hypoxia. Neves et al. demonstrated that QPRT inhibition is associated with impaired differentiation and described a physiological decrease in NAD^+^ levels during differentiation (Neves et al., 2022). In our mature organoids, we observed the association of hypoxia response in the gene sets that were upregulated in the two radial glia and outer radial glia clusters (3/10) as well as with a subset of excitatory neurons (cluster 2). The genes upregulated in radial glia furthermore were associated with glycolysis and gluconeogenesis which supports the hypoxia effect. With respect to the developmental trajectory, we identified an association with mitochondrial organization for genes upregulated as early as 10 days after differentiation start upon QPRT inhibition. This again points towards a potential rescue mechanism of energy preservation and repair mechanisms during stress conditions. In line with this hypothesis, we observed a reduced neuroprotection: The gene *PRDX1* is predominantly expressed in astrocytes and encodes an enzyme protecting cells from oxidative damage by reducing peroxides, including hydrogen peroxide (Guan et al., 2024). We observed a downregulation of *PRDX1* across all cell types, potentially resulting in a diminished redox-signaling capacity. Interestingly, *PRDX1* was downregulated upon QPRT inhibition in all excitatory neuron clusters (Cl1 FDR < 0.001, Cl2/7/11 p < 0.05) but not in any of the inhibitory neuron clusters, specifically highlighting the vulnerability of excitatory neurons to oxidative stress (Papadia et al., 2008). Additionally, PRDX1 was downregulated in a subset of radial glia as well as in astrocytes (p < 0.05). Thus, the combination of a downregulation of neuroprotection with increased hypoxia/oxidative stress and potential accumulation of QUIN might be at the core of the observed effects of QPRT inhibition. We here also identified limited though significant effects on astrocytes: This cell type is specifically involved in regulating neuronal homeostasis, synaptogenesis and the response to metabolic and oxidative stress (Séjourné and Eroglu, 2024). After QPRT inhibition, we observed impaired astrocytic glycolysis which is crucial for the metabolic support of neurons (Zhang et al., 2024). As GAPDH is sensitive to peroxides, this effect is likely linked to and worsened by the downregulation of *PRDX1* as described above. Additionally, *cofilin 1* (*CFL1)* as well as *tubulin alpha 1 a (TUBA1A)* were downregulated in astrocytes of PA-treated organoids. Both are involved in cytoskeleton formation and dynamics (Jiang et al., 2021), i.e., their downregulation potentially impairs astrocyte morphology which in turn impacts on the interactions of astrocytes and neurons.

The gene with the strongest upregulation upon QPRT inhibition at d112 was *EDN1*, a molecular component of the SVZ stem cell niche (Adams et al., 2020). In the brain, *EDN1* plays an important role in the maintenance and proliferation of radial glia and for specific neuronal subtypes (Hostenbach et al., 2016). In addition, it has been reported to increase the number of astrocytes and promote activation under stress conditions such as hypoxia (Hasselblatt et al., 2001). This is consistent with the higher astrocyte proportion observed upon QPRT treatment in our dataset at d112. Adams et al. treated murine neurospheres with EDN1 and found a downregulation of the expression of the proneural transcription factor genes *achaete-scute family BHLH transcription factor 1* (*Ascl*) and *hairy and enhancer of split 6* (*Hes6*, Adams et al., 2020). Interestingly, we also observed a significant downregulation of *HES6* (FDR < 0.001) and *ASCL1* (p < 0.05). Further, Adams and colleagues identified an upregulation of the Notch pathway genes jagged canonical notch ligand 1 (*Jag1)* and hairy and enhancer of split 5 (*Hes5*, which we also observed with nominal significance (Adams et al., 2020). This suggests active Notch signaling in mature organoids upon QPRT inhibition, again pointing towards an ongoing progenitor maintenance to ensure proper neural patterning and maturation (Gonzalez and Reinberg, 2025).

To understand the relevance of our findings in the context of NDDs, we tested for overlapping risk genes. During early organoid differentiation, we report an overlap of downregulated genes with ASD, while at later stages (d80 and d112) we also see overlaps with EP and ID. This points towards a higher vulnerability for the development of ASD at earlier time points of differentiation. In mature organoids, we again find more prominent overlaps with ASD-associated genes across all cells. With respect to cell type-specific effects, we identified strong overlaps of genes for ASD, EP and ID with neuronal subtype DEGs, i.e., cell types developing at maturation stages. Additionally, we find an enrichment in DEGs of several radial glia populations, which further highlights the role of these cell types during neural development. Radial glia, which also showed a strong response to QPRT inhibition, are not only the precursors for the subsequent neuronal differentiation but also support neuronal migration. Interestingly, radial glia dysfunction was also found to be associated with impaired GABAergic neuronal development, similar to our observations in QPRT-inhibited cerebral organoids.

In this study we used PA, a competitive inhibitor of QPRT, which is a small aromatic dicarboxylic acid and might have off-target effects by potentially interfering with other enzymes or metabolic pathways. Further studies must be conducted to compare the effects of QPRT inhibition with QPRT-KO cerebral organoids.

Our single cell sequencing data is limited in that we compared only one organoid per condition. While the comparison across these organoids might be limited in the generalizability due to variability within batches, this dataset allowed us to identify the different subpopulations as direct reference data for the deconvolution. The additionally performed bulk RNA sequencing for 2 independent organoid batches with two replicates per batch and per analyzed time point confirmed the findings from single cell sequencing, as did the immunocytochemistry analysis. Thus, while limited on its own, the scRNA approach in combination with bulk sequencing has proven to be reliable in identifying the reported effects.

## Conclusion

Our study reveals a critical role for the 16p11.2 gene *QPRT* in regulating early human brain development, as modeled in cerebral organoids. Inhibition of QPRT led to significant alterations in organoid morphology, neural progenitor maintenance, and neuronal differentiation, highlighting its importance in maintaining a balanced neurodevelopmental trajectory. The observed reduction in organoid size, neural rosette formation, and excitatory neuron populations, together with the relative increase in inhibitory neurons, indicates a shift towards premature and imbalanced differentiation. These effects are likely mediated by multiple converging mechanisms, including excitotoxicity via QUIN accumulation, disrupted redox homeostasis, metabolic stress, and impaired astrocyte support.

Moreover, transcriptomic changes in QPRT-inhibited organoids show a strong overlap with gene networks implicated in autism spectrum disorder, particularly at early developmental stages, pointing to a heightened vulnerability during this window. Our findings further support the hypothesis that excitation/inhibition imbalance and radial glia dysfunction are shared mechanisms underlying neurodevelopmental disorders.

Taken together, we suggest QPRT as a regulator of early neural development, highlighting its potential involvement in the etiology of neurodevelopmental disorders, especially ASD. It further underscores the utility of stem cell-derived cerebral organoids for modeling gene function during human neurodevelopment, allowing to elucidate cell type-specific effects over time.

## Acknowledgements

We thank Silvia Lindlar for excellent technical support.

This work was funded by “Dr. Elmar und Ellis Reiss Stiftung” and “Freunde und Förderer der Goethe Universität”.

