## Supplementary Figures for "Inhibition of the neurodevelopmental disorder-associated 16p11.2 gene *QPRT* leads to altered cell type distribution in human stem cell-derived cerebral organoids"

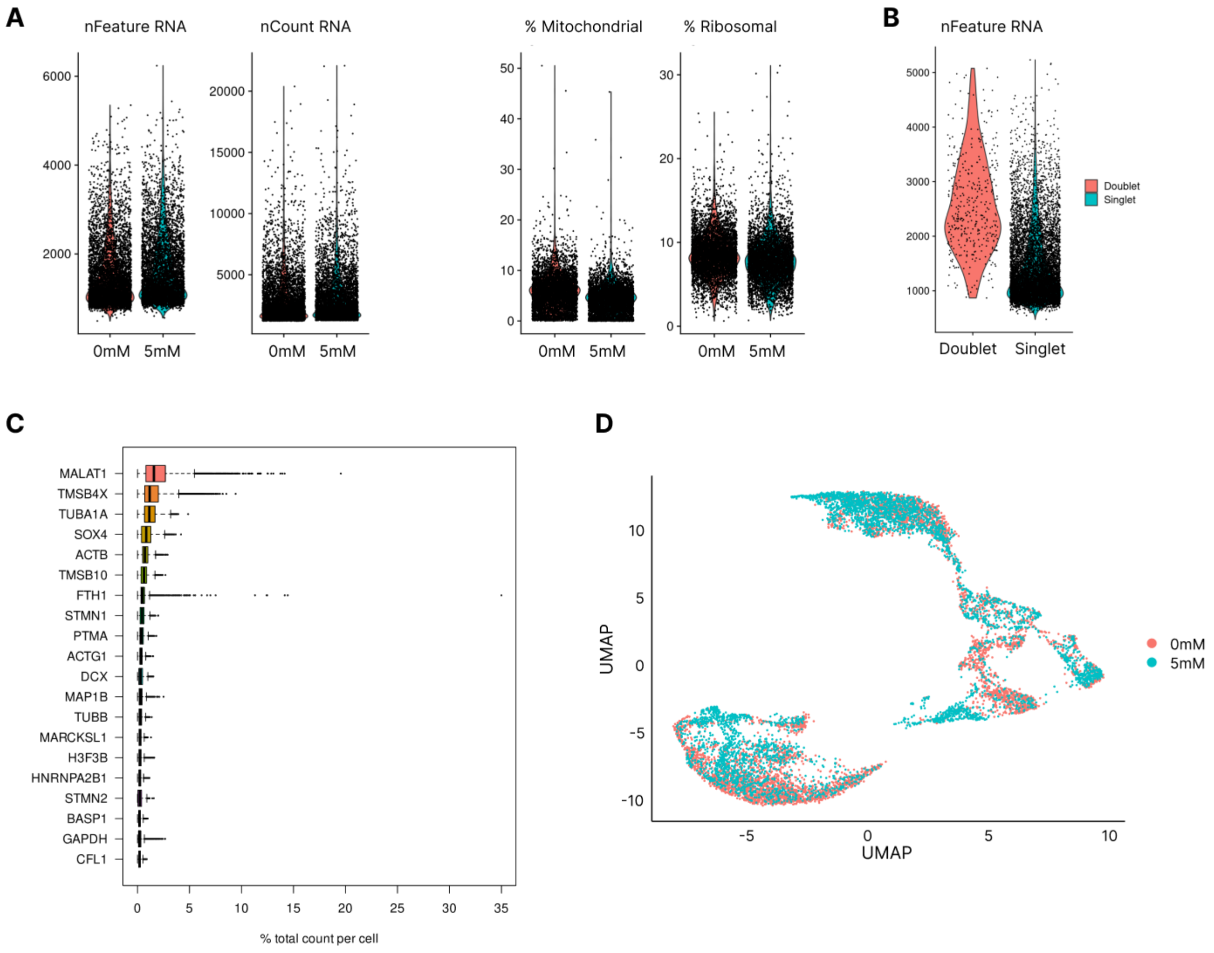


**Supplementary Figure 1.** Quality control of scRNA sequencing data. A) Filtering of low-quality cells. Cells with high percentage of mitochondrial (> 20%) or low ribosomal (< 5%) RNA were filtered out. B) Doublets. Doublet cells (4,16%) were filtered out. C) Highest expressed genes after filtering. The highest expressed genes were housekeeper genes, such as MALAT1 and TUBA1A. D) Clustering after batch correction. UMAP embedding of scRNA data shows insignificant batch effects.

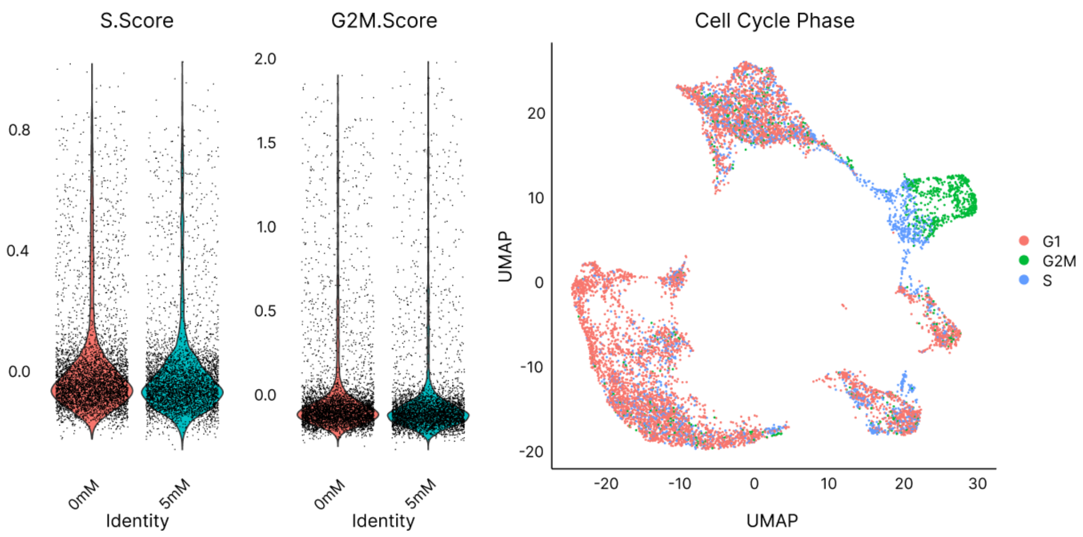


**Supplementary Figure 2.** Cell Cycle Scoring. Violin plot (left) and UMAP embedding (right) of the analysed cells and their assigned cell cycle phase.

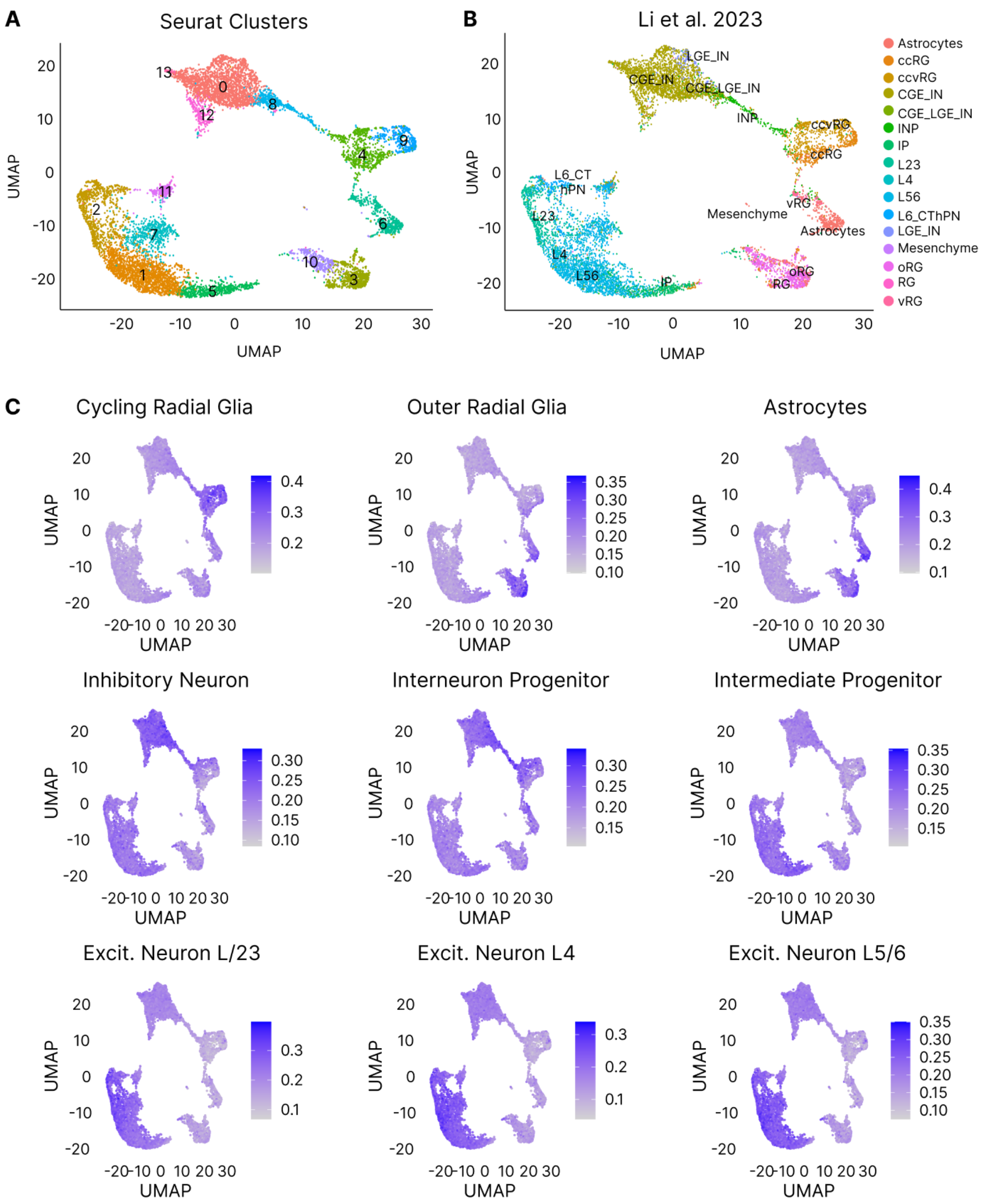


**Supplementary Figure 3.** Clustering. A) Seurat Clusters generated via unsupervised clustering after quality control. Both samples were pooled, clustered at a resolution of 0.6 and plotted in a UMAP embedding, resulting in 14 detected clusters. B) Labelling of cell types using the reference data set from Li et al. ccRG: cycling radial glia; ccvRG: cycling ventral radial glia; oRG: outer radial glia; CGE: Caudal ganglionic eminence; LGE: Lateral ganglionic eminence; IN: Interneuron; INP: Interneuron Progenitor; IP: Intermediate Progenitor; CThPN: cortical thalamic projection neuron C) Probability score for single cell types. Reference Li et al. Color indicates the match safety.

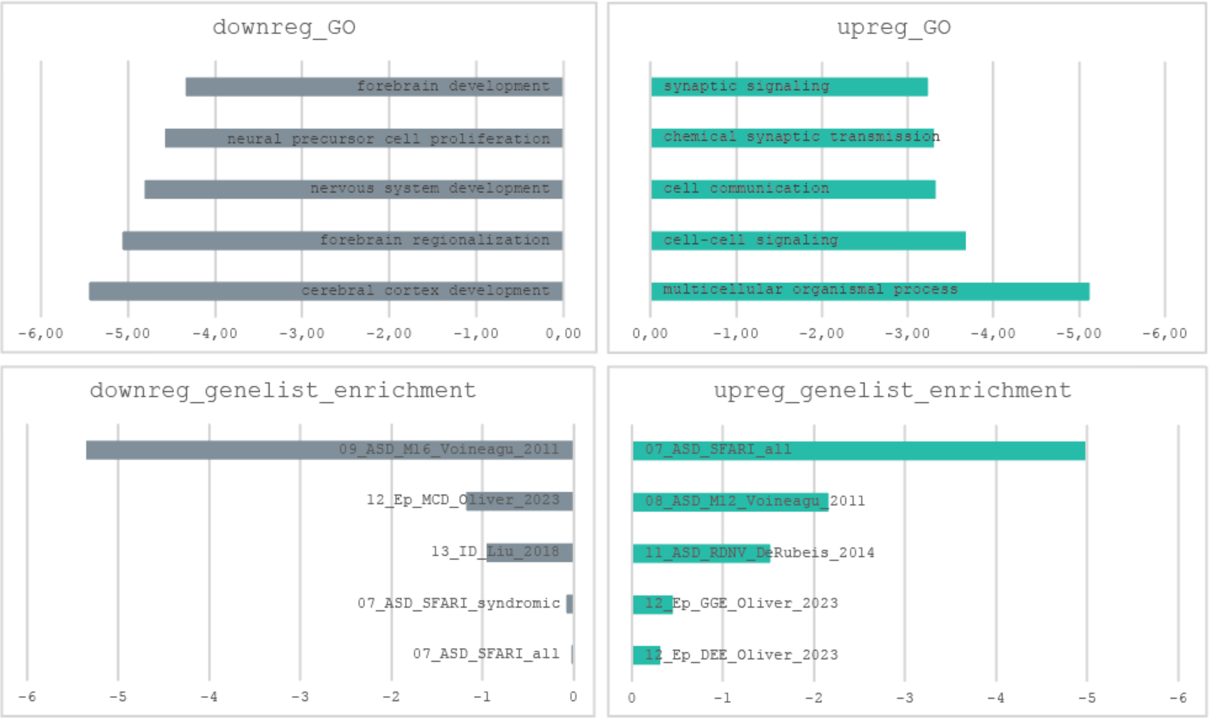


**Supplementary Figure 4.** Enrichment of genes differentially expressed in QPRT-inhibited cerebral organoids at d112 across all cell types for GO terms as well as for disease gene lists.
