## Supplementary Methods for "Inhibition of the neurodevelopmental disorder-associated 16p11.2 gene *QPRT* leads to altered cell type distribution in human stem cell-derived cerebral organoids"

Stem cell maintenance

H9 wild-type (WT) stem cells were maintained on plates coated with Corning® Matrigel® hESC-qualified Matrix (354277) in Essential 8 Flex medium at 37 °C with 5% $CO_2$ in a humidified incubator.

Generation of cerebral organoids and PA treatment

Cerebral organoids were generated using a modified protocol from Esk et al., 2020, which is based on the protocol by Lancaster et al., 2013 (Lancaster et al., 2013; Esk et al., 2020). Two batches were generated following the same procedure (Batch A and B). Briefly, the H9 WT cell line (WiCell WA09) was used to generate cerebral organoids. Three hours before seeding, the medium was replaced with E8 Flex supplemented with 1× RevitaCell (Gibco) to improve cell survival. Cells were dissociated into single cells using Accutase and seeded at 2,500 cells per well in 150 µl of E8 Flex + 1× RevitaCell in low-attachment U-bottom 96-well plates (BIOFLOAT, Sarstedt). Plates were centrifuged at 300 g for 2 minutes and incubated overnight at 37 °C to allow embryoid body formation. On day 3, the medium was replaced with E8 Flex without RevitaCell. By day 5, when embryoid bodies reached ~500 µm with smooth, round edges, the medium was switched to Neural Induction Medium (NIM; DMEM/F-12, 1× N2, 1× Glutamax, 1× MEM-NEAA, 1 µg/ml Heparin) as described in Esk et al., 2020 (Esk et al., 2020). After 10 days, when a bright neuroepithelial layer had formed around a darker core, embryoid bodies were embedded in 15 µl droplets of cold Matrigel on parafilm to promote neural rosette growth. Following a 3-minute incubation at room temperature and 15-20 minutes at 37 °C to allow polymerization, the embedded organoids were transferred into 6 cm dishes containing NIM. On day 13, the medium was changed to Improved NeuroDMEM-A (IDM-A; a 1:1 mix of DMEM/F-12 and Neurobasal, supplemented with 0.5× N2, 1× B27 without vitamin A, insulin, 2-mercaptoethanol, Glutamax, 0.5× MEM-NEAA, and penicillin-streptomycin) as described by Esk et al., 2020 (Esk et al., 2020). During the first two medium changes on days 13 and 14, 3 µM CHIR99021 (GSK-3 inhibitor) was freshly added. Subsequent medium changes up to day 25 were performed without CHIR99021. On day 20, organoid cultures were placed on a shaker at 60 rpm at 37 °C. From day 25, the medium was switched to Improved NeuroDMEM+A (IDM+A; DMEM/F-12 and Neurobasal 1:1 with additional supplements including sodium bicarbonate, N2, B27 with vitamin A, insulin, 2-mercaptoethanol, Glutamax, MEM-NEAA, penicillin-streptomycin, and vitamin C) following Esk et al., 2020 (Esk et al., 2020). Phthalic acid

(PA) was added starting on day 3 to chemically inhibit QPRT. PA was dissolved directly in the respective media (E8 Flex, NIM, IDM-A, or IDM+A), and the pH was adjusted to match control medium.

Organoid Dissociation for Bulk Sequencing

For bulk RNA sequencing, two cerebral organoids per batch and condition were collected for each timepoint. Larger organoids were broken down into smaller cell clusters or single cells using papain. Papain powder (Merck) was dissolved in an activation solution containing 1.1 mM EDTA, 0.067 mM 2-mercaptoethanol, and 5.5 mM L-cysteine hydrochloride to a concentration of 250 U/ml, then activated at 37 °C for 30 minutes. The dissociation solution was freshly prepared by combining activated papain (final 20 U/ml), DNase I (final 125 U/ml, Merck), and Hank's Balanced Salt Solution (HBSS, Life Tech). Organoids were minced with pipette tips to aid digestion and incubated at 37 °C on a shaker at 150 rpm. After 30 minutes, organoids were further triturated to reduce clumping. Additional incubation and trituration were applied, if necessary, until most clumps were removed. Dissociated cells were pelleted and stored at - 80 °C until RNA extraction.

**Code for image analysis**

Image analysis was done using Fiji (Version 1.54i, (Schindelin et al., 2012) and the following code (with use of Open AI, September 2024):

```
// Select directory with images
dir = getDirectory("Choose a directory containing TIFF images");
list = getFileList(dir);
// Iterate through all images in the directory
for (i = 0; i < list.length; i++) {
    if (endsWith(list[i], ".tiff")) {
        // Open image
        open(dir + list[i]);
        // Duplicate the image and split channels
        run("Duplicate...", "title=Duplicate duplicate");
        run("Split Channels");
        // Set threshold, create mask, and fill holes for Channel 1
        selectImage("C1-Duplicate");
        setThreshold(771, 65535, "raw");
        run("Create Mask");
        resetThreshold();
        run("Fill Holes");
        // Perform particle analysis
        run("Analyze Particles...", "size=300000-Infinity display clear add");
        // Measure ROI for all channels
        for (j = 1; j <= 4; j++) {
            selectImage("C" + j + "-Duplicate");
            roiManager("Select", 0);
            roiManager("Measure");
        }
        // Save results as CSV
        saveAs("Results", dir + "Measurements_" + list[i] + "_Results.csv");
        // Close image and masks
        close("*");
        run("Clear Results");
    }
}
```
